# The Supplementary Eye Field Tracks Cognitive Efforts

**DOI:** 10.1101/2021.01.14.426722

**Authors:** Julien Claron, Julie Royo, Fabrice Arcizet, Thomas Deffieux, Mickael Tanter, Pierre Pouget

**Author notes:** These authors jointly supervised the work (Mickael Tanter, Pierre Pouget).

## Abstract

Pupil dilation is known to be an index of cognitive effort. Nevertheless, our lack of knowledge of the precise dynamics through which pupil size and activity of the medial prefrontal cortex are conjugated during cognitive tasks highlights the need for its further investigation before, during, and after changes in pupil size. Here, we tested whether pupil dynamics are related to the activity of the supplementary eye field (SEF) during a mixed pro/anti-saccade oculomotor task in two macaque monkeys. We used functional ultrasound imaging (fUS) to examine temporal changes in brain activity at the 0.1-s time scale and 0.1-mm spatial resolution in relation to behavioral performance and pupil dynamics. By combining the pupil signals and real-time imaging of NHP during cognitive tasks, we were able to infer localized CBV responses within a restricted part of the dorsomedial prefrontal cortex, referred to as the SEF, an area in which anti-saccade preparation activity is also recorded. Inversely, SEF neurovascular activity measured by fUS imaging was found to be a robust predictor of specific variations in pupil diameter over short and long time scales. Furthermore, we directly manipulated pupil diameter and CBV in the SEF using reward and cognitive effort. These results demonstrate that the SEF is an underestimated but pivotal cortical area for the monitoring and implementation of cognitive effort signals.

---

Seminal studies revealed that pupil dilation varies with increasing task demands, including perception, attention, task consolidation, learning, and memory^1–6^. Two dominant interpretations for these findings have been proposed. Numerous authors concluded that pupil dilation reflects the demands of a task, whereas others took it a step further and proposed that pupil dilation actually reflects the effort exerted in response to such demands^1,2,7^. The precise neural substrates by which such cognitive processes influence pupil diameter are still unclear, but inputs from the dorsal part of the medial prefrontal cortex (dmPFC), which mediates arousal, are likely involved.

The dmPFC contains the frontal eye fields (FEF), supplementary motor area (SMA), and supplementary eye field (SEF). The FEF is known to be involved in the control of eye movements and attention and recent studies have shown that the amplitude of pupil responses depends on the combination of the light stimulus and subthreshold FEF electrical microstimulation^8,9^. Strongly interconnected to the FEF and anterior cingulate cortex (ACC), the SEF is a key region that integrates attentional, short memory, and oculomotor tasks^10,11^. The SEF also directly projects to the brainstem oculomotor nucleus. In principle, the ACC, SEF, and FEF networks may directly modulate the olivary pretectal nucleus, which encodes retinal illumination and directly activates the pupil-constrictor pathway. Alternatively, or in addition, the ACC, SEF, and ACC networks may act indirectly through the occipital visual cortical areas, in which the visual responses are modulated by FEF and may, in addition to programming the oculomotor plan, participate in the pupil light reflex (PLR). Although the function of the SEF in oculomotor tasks is reasonably well defined, the relationship between SEF activity and pupil dynamics is still unknown.

Pupil dynamics have been studied during preparation and before the execution of eye movements during oculomotor protocols^12^, suggesting valid qualities for ongoing cortical processes. In the context of the anti-saccade task, subjects are instructed before the appearance of a stimulus to either automatically look at the peripheral stimulus (pro-saccade) or suppress the automatic response and voluntarily look in the opposite direction from the stimulus (anti-saccade). In this type of paradigm, pupil size was found to be larger in preparation for correct anti-saccades than in preparation for correct pro-saccades and erroneous pro-saccades made in the anti-saccade condition^13^. When an incorrect saccade is executed with latencies in the range of express saccades, execution of the movement indicates that subjects are unable to inhibit involuntary actions, whereas they have no difficulties in generating voluntary saccades if they correct such directional errors. Given that the SEF is known to be critically involved in the production of antisaccades^14^, the precise dynamics through which pupil size and SEF activities are conjugated merits further investigation before, during, and after pupil size modulation.

This question can be addressed using modern neuroimaging techniques, such as functional ultrasound (FUS) imaging. This innovative imaging technique allows very precise mapping of fine temporal changes in brain neurovascular activity at high spatial resolution in large cortical areas in non-human primates^15,16^. In the present series of experiments, we tested whether pupil dynamics are linked to SEF activity during an anti-saccade task on awake monkeys.

We obtained two primary results: 1) SEF activity is a robust predictor of specific variations of pupil diameter over both short (milliseconds) and long (minutes) time scales and 2) strong covariations of pupil diameter and CBV can be selectively observed in the SEF by manipulating reward and cognitive effort.

## RESULTS

We recorded SEF activity by fUS imaging in two monkeys (n = 26 sessions for Monkey S and n = 20 sessions for Monkey G) trained to perform a pro/anti-saccade task (Fig.1.a). Both monkeys performed the task reliably across all recording sessions and the average correct rate of both monkeys was approximately 85% (Figs.1.b and 1.c). The two monkeys showed significant shorter latencies for pro-saccades than antisaccades, confirming a higher cognitive effort when an anti-saccade was planned (Monkey S: 197 ± 15 ms for pro-saccades, 262 ± 10 ms for anti-saccades, *p* = 7e-9; Monkey G: 218 ± 32 ms for pro-saccades, 267 ± 24 for anti-saccades, *p* = 2e-4, using Wilcoxon’s rank test). All trials were used for our analysis regardless of their nature (pro-saccades versus anti-saccades, right vs. left) or whether they were successful.

**Figure 1.**
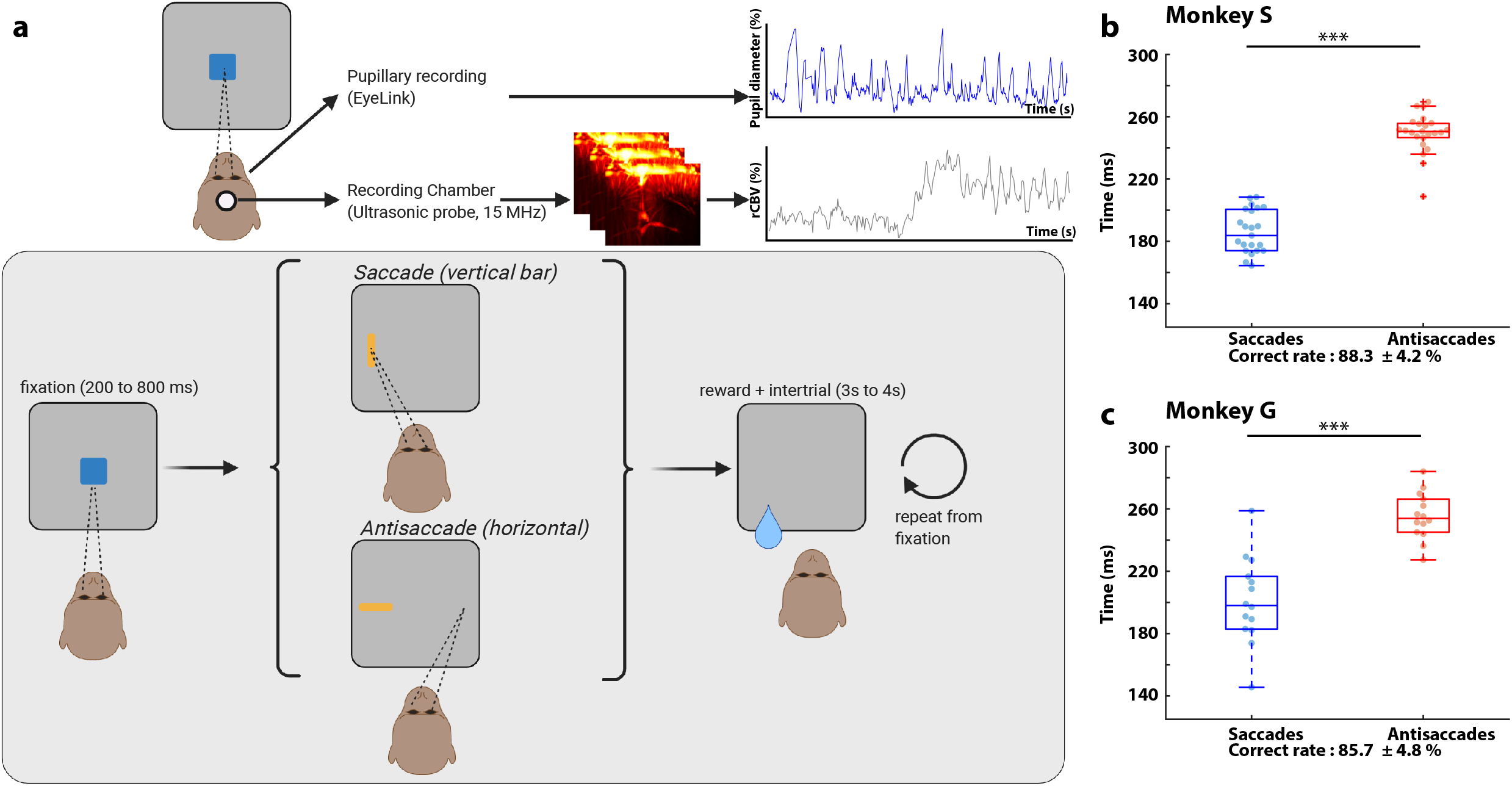
Task timeline. (a) The heads of the monkeys are fixed in a chair with a 15-MHz ultrasonic probe in a recording chamber. An EyeLink recording system records the eye position and pupillary diameter in real time. During the task, there is a baseline of 200 to 220 s (randomized) and a fixation point is shown. If the animal succeeds, a pro-saccade (vertical rectangle) or an anti-saccade (horizontal rectangle) is shown on the screen. Based on the hint, the animal performs a saccade or an anti-saccade and, if he succeeds, receives a reward associated with a specific color of the fixation point (red: 0.5 times the normal reward, blue: 1 time the normal reward, green 1.5 times the normal reward). This action is followed by a grey screen of 3 to 4 s (randomized) used as an intertrial before repeating from the fixation point. (b) The average saccade response time for Monkey S for all sessions was 197.0 ± 15 ms and the anti-saccade response time 262.0 ± 10 ms, with a total correct rate of 88.2 ± 4.2%. (c) The average saccade response time for Monkey G was 218 ± 32 ms, and the antisaccade response time 267 ± 24 ms, with a correct rate of 85.7 ± 4.6%.

### 1 – Determination of the functionally activated area without choosing *a priori*

We wished to investigate the relationship between pupil size and brain activity without any *a priori* choice concerning the activated area. We thus applied the generalized linear model to the fUS data using pupil diameter as the input matrix. In total, 600 trials were used for these analyses and the time between each trial was a random interval of 3 to 4 s determined after each trial. We constructed the input matrix by realigning all pupil diameters with the target presentation time and took the value of the pupil diameter at the first local maxima. Indeed, the pupil diameter was higher for the first trials (in blue in Figs. 2.a and 2.d) than for the last (in red in Figs. 2.a and 2.d). We compensated for this observed decrease in pupil diameter during the trials by using the pupil diameter at maximum dilation (at 0.8 s for monkey S, (Fig 2.a) and 0.6 s for monkey G. (Fig 2.d.). The highlighted pixels in Fig 2.b and Fig 2.e are those for which p < 0.05 (before Bonferroni correction), indicating that the pixels highly correlated with the pupil diameter in our cortical imaging plane. In these activated pixels, mostly consisting of the surface of the cortex, we found the activated area to be in the SEF, bilaterally for monkey S and mostly in the left area for monkey G. Such activation is consistent with the Paxinos atlas for the localization of the SEF. High correlations of the fUS signals with the pupil diameter were found in the SEF regions for both animals. Finally, we extracted the cerebral blood volume (CBV) temporal signal from our fUS data by spatially averaging the signal isolated in the functionally activated area (Figs. 2.c. and 2.f). The co-variation between CBV and pupil signals occurred over short time scales, as the fUS signal exhibited a large and sharp increase in CBV in response to the single first trial (zoom in Figs. 2c and 2f) of each task. This sharp increase in CBV is consistent with the results of previous studies^15^.

**Figure 2.**
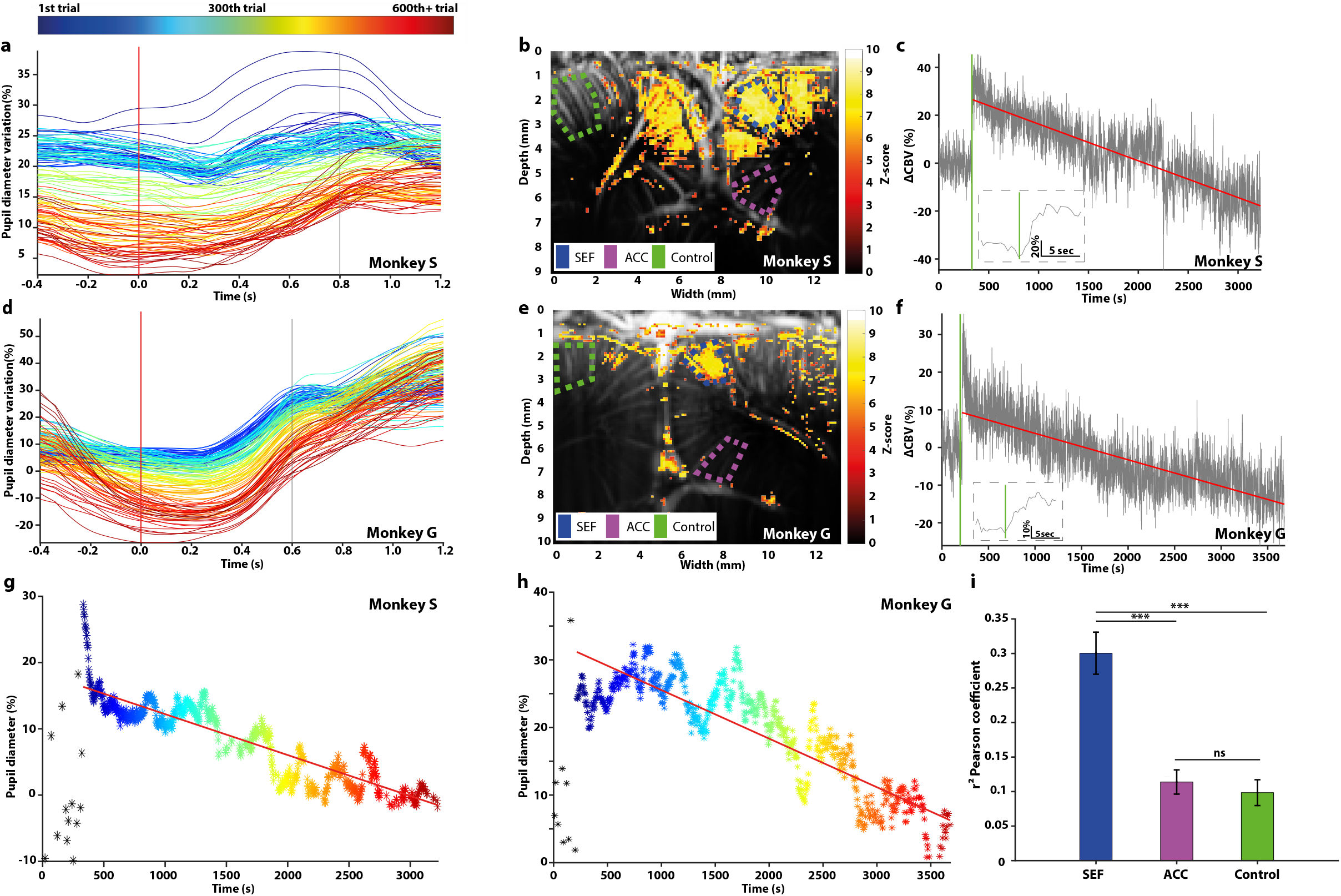
Example of one session for each monkey of the vascular and pupillary responses. (a) Pupillary response over time. The colour represents the number of the trial (blue: 1st trial, red: last trial), whereas the temporal abscissa represents the time prior to the presentation of the saccade or anti-saccade hint. The vertical line represents the chosen time for the maximal dilation (0.8 s for Monkey S). (b) The vascular response of Monkey S using Fig. 2.g as an input matrix for the generalized linear model (GLM). The background image consists of an anatomical image obtained by averaging all the Doppler films. The Z-score map was obtained using the GLM and thresholded using the Bonferroni correction (p < 0.05 uncorrected). (c) Example of a CBV response during the starting of a task for Monkey S., showing a step at the end of the baseline and the beginning of the task. The small square represents a zoom on the 1^st^ trial. (d) Same as for 2.a for Monkey G. The vertical line is at 0.6 s after presentation of the hint. (e) Same as for 2.b for Monkey G. using 2.h as an input matrix (f) Same as for 2.c for Monkey G. (g) Maximum pupillary dilation at 0.8 s after presentation of the hint using the same colour code as in B. Black stars represent the pupillary diameter during baseline. (h) Same as for 2.g for monkey G. (i) Pearson’s correlation coefficient between pupil diameter (see h and j) and three areas in the brain: SEF (blue), ACC (purple), and Control (green) for all non-reward-modulated sessions for both animals (n = 21 sessions for Monkey S, n = 13 sessions for Monkey G), ***p < 0.001, ns: not significant. All vertical green bars represent the end of the baseline and the start of the task.

### 2 – Long-term covariation of CBV in the SEF and pupil diameter

During the successive trials of a non-reward-modulated pro-saccade and anti-saccade task, we observed a large and reproducible decrease in the relative CBV (rCBV) of the SEF, defined anatomically using the monkey brain atlas and functionally using the GLM, as previously described (Figs. 2.c and 2.f) (−23 ± 2%/h for Monkey S and −21 ± 5%/h for Monkey G) after the initial step^15^. Given the strong correlation between pupil size and cognitive engagement in the task, we examined the pupil diameter after a pro-saccade or anti-saccade task. We also observed a large and reproducible decrease in pupil diameter throughout the duration of the session (Figs. 2.g and 2.h, - 9.3 ± 0.3%/h for Monkey S and −9.7 ± 3.1%/h for Monkey G) on a long time scale. The decreases in pupil size correlated with the change in activity of the SEF. Furthermore, we compared the squared Pearson’s r^2^ correlation coefficient between three regions of the brain (SEF, ACC, and a control area, anatomically chosen for the SEF and ACC and to be the non-activated cortical area farthest from the SEF) and the pupil signal. The r^2^ was significantly higher (using a linear mixed-effects statistical model) in the SEF (0.30 ± 0.03, sem) than in the ACC (0.11 ± 0.02, sem) or control area (0.10 ± 0.02) (Fig. 2.i). Such a correlation between the activity in the SEF and pupil diameter has not yet been reported.

### 3 – Timing of the SEF and pupil activities

First, we determined whether SEF activity and pupil dynamics were simultaneous or whether one activity occurred before the other. This required that we first determine the hemodynamic response function (HRF) describing the neurovascular coupling for each of the monkeys. We thus averaged the activity in the SEF area during a task and fitted it with a gamma-inverse density function for each primate to establish the HRF for each (n = 9 sessions for Monkey S and n = 7 sessions for Monkey G, choosen over the first few sessions. Sessions were kept if the non-linear fitting algorithm managed to converge in less than 400 iterations). We obtained the parameters and individual fits by session and plotted the average (Figs. 3.a and 3.d). We performed the same type of experiment to determine the pupillary light response function (PLR) (represented in Figs. 3.b and 3.f) in which the minimum of the function represents the lag of the pupil response (440 ms for Monkey S and 480 ms for Monkey G). For example, we superimposed the pupil (Figs. 3.c and 3.g; red) and hemodynamic (Fig. 3.c. and 3.g; grey) responses for one session to search for a shift in the SEF hemodynamics and pupil activities. Comparison between the histogram of each fUS peak for each trial, minus the peak of the HRF, and the histogram for pupil peak for each trial, minus the peak of the PLR for the pupil, showed a lag between the CBV and pupil response (Figs. 3.d and 3.h), suggesting that the two events do not occur at the same time in the brain. On average, the SEF responses were found to precede the pupil responses.

**Figure 3.**
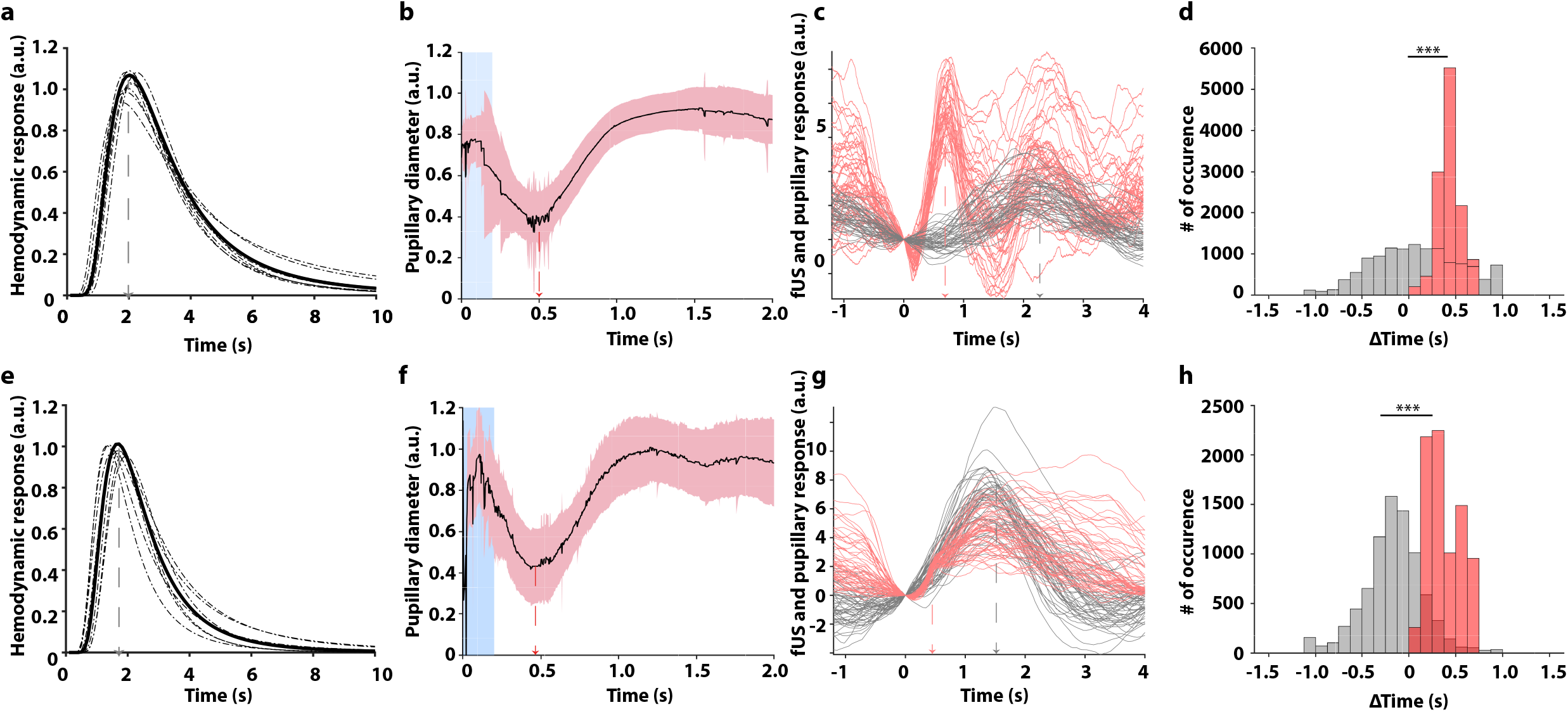
Pupillary and fUS response and comparison of the lag. (a) Hemodynamic response function of Monkey S. The maximum peak occurred at 2.0 s. (b) Pupillary light response of Monkey S. The peak occurred at 0.44 s. The blue rectangle represents the duration of the white flash. (c) Pupillary response (pink) and superimposed vascular response (grey) for each trial. All the data were strictly zeroed at t = 0 s. (d) Histogram of pupillary response (red) and fUS response (grey) prior to execution of the saccade minus the max of the HRF and PLR. (e-h) same for Monkey G. The HRF peaked at 1.7 s and the PLR at 0.48 s. ***p < 0.001 (Wilcoxon’s rank test)

Then, we directly modulated the level of reward and cross-compared the reward levels with the cognitive effort of each trial to manipulate task engagement and further examine the relationship between variations in pupil diameter and SEF activity. Modulation of the reward was reflected in both changes in pupil diameter and SEF activity. We next examined the association between cognitive effort and SEF activity.

### 4 - Reward magnitude and task difficulty modulate SEF activity and pupil diameter

We then measured engagement in the task by slightly adapting our paradigm by adding a color code at the fixation point on each given block of 100 trials and modifying the magnitude of the reward. The potential reward delivered for each correct pro- or antisaccade was red for 0.5 reward units, blue for 1.0 reward unit, and green for 1.5 reward units (Figs. 4.a and 4.b). During the task, the ΔCBV changed during the transition from one reward level to another, as did pupil dilation. We also observed a slight disengagement of primates when the task is high-cognitive demanding (e.g. antisaccade) for a low reward (e.g. 0.5 reward unit for red fixation point).

**Figure 4.**
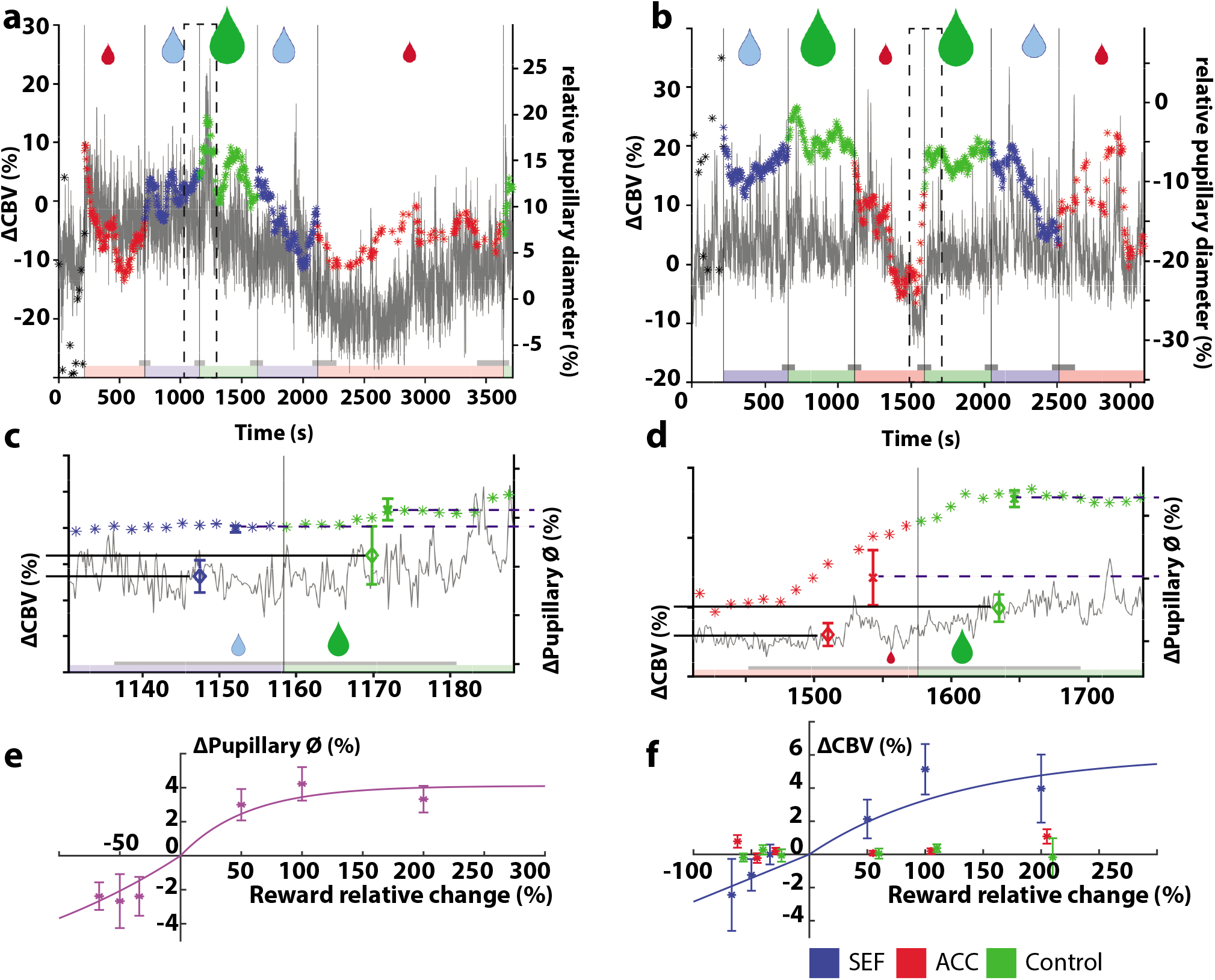
Reward modulation during the task for both Monkey S and Monkey G. (a) CBV and pupillary response over one session for Monkey S. The colours represent the quantity of reward obtained after a successful trial (green: 1.5X the base reward, blue: 1X the base reward, and red: 0.5X the base reward). (b) Same as for (a) for Monkey G. (c) Zoom of (a) for one reward transition at approximately 2,500 s. The grey area under the curve represents the last 10 trials before transition and the first 10 trials after transition, which were used to calculate the CBV and pupillary values before and after the transition (diamond for CBV, star for pupil). (d) Same as (c) for Monkey G. (e) Pupil dilation according to the change in reward for monkeys S and G. Mean ± standard error of the mean. (f) The CBV changed according to the change in reward, all sessions for Monkeys S and G. Blue represent the SEF, red the ACC, and green the control area. Mean ± standard error of the mean.

We wanted to quantify pupil dilation and the ΔCBV during such transitions. We computed the transition between the two levels of reward by computing the average CBV and pupil diameter for 10 trials before the transition and 10 trials after. The difference between the values after versus before the transition gives the increase or decrease induced by the transition (Fig. 4.c for Monkey S and Fig. 4.d for Monkey G). Pupil dilation decreased in the transition from a higher to lower reward and increased in the transition from a lower to higher reward (Fig. 4.e). We observed an increase in the ΔCBV in the SEF during a transition from a lower to higher reward, but no statistically significant measure was obtained for a transition from higher to lower reward (Fig. 4.f., in blue)

Interestingly, we observed a marginal effect on the ACC (Fig. 4.f, in red) for the reward transition from 0.5 units to 1.5 units and 1.5 units to 0.5 units. More rostral investigations of the ACC would be required to measure such effects in this area. A control area showed no augmentation or decrease (Fig. 4.f, in green). Finally, increasing the reward resulted in augmentation of both pupil dilation (Fig. 4.e) and ΔCBV in the SEF (Fig. 4.f, blue). Overall, the fUS activity in the SEF was strongly modulated in real-time by the reward, as was pupil size, showing that the SEF may play a role in the cognitive efforts for such demanding oculomotor tasks.

## DISCUSSION

We combined pupil signals and real-time imaging of NHP during cognitive tasks, which allowed us to infer localized CBV responses within a restricted part of the dorsomedial prefrontal cortex, referred to as the SEF, an area in which anti-saccade preparation activity is also recorded. Inversely, SEF neurovascular activity measured by fUS imaging was found to be a robust predictor of specific variations in pupil diameter over short and long time scales. The manipulation of reward and cognitive efforts performed by the animals resulted in strong temporal covariation of pupil diameter and CBV within the SEF. Overall, these results show the region of the SEF to be an underestimated pivotal element within the medial frontal cortex of primates for monitoring and implementing the cognitive effort signals observed within autonomous networks.

In previous studies, SEF neurons have been shown to participate in the selection of eye movements by representing the context-dependent action value of various possible oculomotor behaviors^17^. However, the SEF alone does not have the aptitude to directly select eye movements^17^. In the same vein, the SEF does not directly participate in the rapid inhibitory control of eye movements in response to sudden changes in task requirements. Our results showing covariations of pupil diameter and CBV within the SEF have important implications because variations in pupil diameter have been observed for a variety of tasks^1–5^. Two principal explanations have been provided to account for such pupil-effort covariation. First, a direct “bottom-up” influence on decisions to produce a bias towards accepting an effort. This would be consistent with the widely held view that the strength of neural representations for choice attributes directly influence the decision. For example, it has been shown that intensifying encoded rewards through the simulation of future episodic events is linked with decisions that promote higher long-term payoffs and even increase prosocial behavior.

As for neural implementation, phasic locus coeruleus (LC) activity is known to transmit feedforward information to the SEF via ascending projections to the prefrontal cortex (PFC), providing a plausible pathway for such a bottom-up influence. Neural readout of the autonomous activation associated with arousal could provide an additional mechanism by which the arousal signal observed here may bias choices, serving as a signal that the organism is indeed ready to accept the physical challenge.

In the ACC, unlike the SEF, there was not even a tendency of heightened CBV modulation under conditions of cognitive effort. This finding is compatible with an earlier report showing that ACC neurons in the monkey are not selectively active during the countermanding of saccades, an operation assumed to involve cognitive effort and inhibition of action^18^. However, it stands in sharp contrast to a large body of literature, based on functional MRI imaging in humans, indicating that activation in the ACC is strongly heightened under conditions of effort^19–22^.

There are several possible explanations for this discrepancy. There may be a species-specific difference, such that neurons in the human ACC monitor cognitive effort, whereas those in the monkey ACC do not. This seems improbable because, in general, anatomically homologous areas appear to serve similar functions in the two species^23^. This cannot, however, be altogether ruled out. The human ACC possesses a cell type not found in the monkey ACC^24^ and, therefore, may serve a function not served by the monkey ACC. It is possible that our recording sites lay outside the region of the ACC responsible for effort-related activity^25^. This also seems improbable because we deliberately recorded in the subregion that is connected to the SEF^26^ and in which, accordingly, it would be most reasonable to expect to find activity sensitive to cognitive effort in an oculomotor task. It is also possible that the cognitive-specific bold signals detected in human fMRI studies are related to neural events other than spiking activity and CBV, for example, presynaptic potentials^27^. It may be the case that ACC activity, even in humans, does not exhibit enhanced spiking activity under conditions of cognitive effort. For the ACC to serve a cognitive effort and alert the rest of the cortex to the presence of cognitive effort would require enhanced spiking activity because spikes are the currency used between the ACC and other cortical areas. Thus, the remaining conclusion is that the ACC does not monitor cognitive effort.

Overall, our observations are consistent with a possible top-down influence from the SEF to the noradrenaline arousal system, which may serve to transmit information about the commitment to overcome a great physical demand, thus resulting in automatic accelerating upregulation of arousal states to prepare the organism for the upcoming challenge associated with the recent choice. As SEF activity precedes pupil modulation, our results also allow us to conclude that within the medial frontal cortex of primates, aside from the ACC, the SEF may also play a pivotal role in the implementation of the cognitive effort signals observed within autonomous networks.

## Acknowledgments

We thank Jeffrey Schall and Jérôme Sallet for helpful comments on earlier versions of the manuscript. This work was funded by a research grant from the European Research Council (ERC) under the European Union’s Seventh Framework Program (FP7/2007–2013)/ERC Advanced grant agreement no. 339244-FUSIMAGINE and the Fondation Bettencourt-Schueller. It was also supported by the Inserm Accelerator of Technological Research in Biomedical Ultrasound.

## METHODS

### Animal model and behavioral data

All experiments were ethically approved by the French “Ministère de l’Education, de l’Enseignement Supérieur et de la Recherche” under the project reference APAFIS #6355-2016080911065046. Functional data were acquired from two captive-born rhesus monkeys (*Macaca mulatta*), S and G, trained to perform various types of visual tasks. In the saccade task, the animal has to fix its gaze on the cue object presented on the right or left side of the screen; in the antisaccade task, it has to fix its gaze on the opposite side from where the cue appeared. Each animal performed at baseline (200 to 220 s, random) followed by saccades and anti-saccades (randomized) over 1 h. During data acquisition, the eye position of the primate was monitored at 1 kHz using an infrared video eye tracker (Eyelink 1k, SR-Research), which enabled live control of the behavioral paradigm and the delivery of a reward based on the success or failure of a visual task^28^.

### Experimental setup

We recorded 46 sessions (26 for Monkey S and 20 for Monkey G) of pro-saccade and anti-saccade tasks, with two kinds of sessions (Fig. 1).

The conventional session, without reward modulation, consisted of only a blue square before the pro-saccade or anti-saccade cue and the reward was kept constant within and between all sessions. For the second type of task, with reward modulation, the same basal reward was retained and the animal was presented with three colored dots (red for 0.5 reward unit, blue for 1 reward unit, and green for 1.5 reward unit) before the pro-saccade or anti-saccade cue. Behavioral data, such as pupil diameter, were recorded with an EyeLink system and cerebral blood volume (CBV) using a functional ultrasound scanner for all sessions.

All tasks were driven by EventIDE software (OkazoLab, Netherlands).

The reward was calibrated to the weight of the primate and the model of the rewarding tube (approximately 30 ms/kg for the electronic valve), which delivered sugary water. Primates were under mild fluid restriction (approximately 30 mL/kg/day) and could drink *ad libitum* while working.

### Implant and probe for functional ultrasound imaging for awake cooperative monkeys

The head of the monkey was fixed using a surgically implanted titanium head post (Crist Instrument, MD, USA). After behavioral training of the animals, a recording chamber (CILUX chamber, Crist Instrument, MD, USA) was implanted and a craniotomy (diameter 19 mm) was performed (mediolateral: +0 mm, anteroposterior : +26 mm). A custom ultrasonic probe (128 elements, 15 MHz, 100 x 100 μm2 of spatial resolution) with sterile ultrasonic gel was used in the chamber. The acquired images had a pixel size of 100 x 100 μm and a slice thickness of 400 μm.

### Functional ultrasound (fUS) recording

Changes in CBV were measured using a real time functional ultrasound scanner prototype (Iconeus and Inserm U1273, Paris, France) with a 15-MHz linear probe. Data were acquired by emitting continuous groups of 11 planar ultrasonic waves tilted at angles varying from −10° to 10°. Ultrasonic echoes were summed to create a single compound image acquired every 2 ms. Final Doppler images were created by averaging 200 compound ultrasonic images after spatiotemporal filtering based on the singular value decomposition of the ultrasonic images.

### Eye movements and pupil recordings

Eye movements and pupil diameter were recorded during the tasks using a video eye tracker (Eyelink 1k, SR-Research) connected to an analog-to-digital converter (Plexon Inc, TX, USA). All data were collected using Plexon software and analyzed using MATLAB (The MathWorks Inc., Massachusetts, USA). Saccades were detected when the eye’s horizontal velocity went over 30°.s^-1^.

### Data processing

#### Generalized linear model

Doppler data were analyzed using a generalized linear model approach implemented in Matlab. The stimulation pattern in the design matrix was convoluted with the fUS-determined HRF and a Z-score and p-value map were obtained. The activation maps show the Z-score of all pixels in the images with a p-value < 0.05 (before Bonferroni correction). We chose the region of interest (ROI) within the supplementary eye field based on the Z-score map and Paxinos atlas for macaque brains and the signal was averaged to obtain a single temporal signal. The spatially averaged signal was then expressed as the relative increase in CBV (in percent) by subtracting the baseline CBV (calculated during the baseline at the beginning of an acquisition) followed by division of the difference by the baseline CBV.

#### Determination of the pupil diameter

The pupil diameter was expressed in percent by subtracting the baseline value and then dividing the difference by the baseline value (in which we excluded all blinks and moments in which the eyes were closed). We then determined the maximum dilation diameter following a task by realigning the pupil diameter at the onset of the cue presentation. We chose the 1^st^ local maximum of the pupil diameter (0.8 s after target onset for Monkey S and 0.6 s after target onset for Monkey G) to extract the pupil diameter for the i^th^ trial.

#### Fitting of the hemodynamic response

The hemodynamic response was determined by averaging the CBV response of all trials and fitting the average by an inverse-gamma probability distribution the using MATLAB *lsqcurvefit* (Optimization Toolbox) algorithm for least square non-linear fitting, as previously described by other authors^29^.

#### Determination of the pupillary light response

The Pupillary light response (PLR) was determined over 1 single session by averaging the pupil diameter response to a short white flash (200 ms) after fixation (250 ms). The flash was followed by a black screen for 4 s to ensure that the pupil returned to the basal state before the following fixation/flash.

#### Statistical analysis of the hemodynamic responses

Statistical analysis between two groups were performed using the Wilcoxon rank test, due to the non-normality of our data, using the Matlab *ranksum* function, the null hypothesis being no statistical difference between the two groups. If more than two groups were available and the data hierarchically organized, we used a linear mixed statistical model. Data were homogenized using a square root transformation and the variance of homogeneity assessed using the Bartlett test and residual normality the Shapiro-Wilk test.

